# Characterizing the Theory of Mind Network in Schizophrenia Reveals a Sparser Network Structure

**DOI:** 10.1101/2020.06.03.124834

**Authors:** Florian Bitsch, Philipp Berger, Arne Nagels, Irina Falkenberg, Benjamin Straube

**Affiliations:** Department of Psychiatry and Psychotherapy, Philipps-University Marburg, Rudolf-Bultmann Str. 8, 35039 Marburg, Germany; Department of English and Linguistics, Johannes Gutenberg-University Mainz, Jakob-Welder-Weg 18, 55128 Mainz, Germany

**Keywords:** effective connectivity, disconnectivity, mentalizing, psychosis, social cognition

## Abstract

Impaired social functioning is a hallmark of schizophrenia and altered functional integration between distant brain regions are expected to account for signs and symptoms of the disorder. The functional neuroarchitecture of a network relevant for social functioning, the mentalizing network, is however poorly understood. In this study we examined dysfunctions of the mentalizing network in patients with schizophrenia compared to healthy controls via dynamic causal modelling and an interactive social decision-making game. Network characteristics were analyzed on a single subject basis whereas graph theoretic metrics such as in-degree, out-degree and edge-connectivity per network node were compared between the groups. The results point to a sparser network structure in patients with schizophrenia and highlight the dorsomedial prefrontal cortex as a disconnected network hub receiving significantly less input from other brain regions in the network. Further analyses suggest that integrating pathways from the right and the left temporo-parietal junction into the dorsomedial prefrontal cortex were less frequently found in patients with schizophrenia. Brain and behavior analyses further suggest that the connectivity-intactness within the entire network is associated with functional interpersonal behavior during the task. Thus, the neurobiological alterations within the mentalizing network in patients with schizophrenia point to a specific integration deficit between core brain regions underlying the generation of higher-order representations and thereby provide a potential treatment target.

## 1. Introduction

Discovering the pathophysiology of psychiatric disorders by examining communication deficits between distant brain regions is a promising trend in clinical neuroscience [1,2]. This perspective seems particularly appropriate to understand the psychopathology of schizophrenia, given that the symptoms of the disorder have been interpreted as signs of disconnections between neural circuits [3]. More than 100 years later, this idea still receives support [4,5,6]. Within this framework it is assumed that a dysfunctional integration between functionally specialized brain systems is linked with deficits in forming learning dependent representations. These are assumed to extend to the highest form of mental representations, the mental states of others [7]. These theoretical assumptions are underlined by recent empirical evidence pointing to altered functional [8,9,10,11,12] and structural network architecture in patients with schizophrenia [8,11,13]. Along these lines, the probability of alterations in a brain region increases with the region’s interconnectedness to other brain regions. This led to the assumption that the connectivity-rich frontal and temporal hubs are particularly vulnerable in schizophrenia [13,14].

A hallmark of schizophrenia is impaired social functioning [15] which has been associated with dysfunctional mentalizing abilities, i.e. the inference and representation of other people’s mental states [16]. Alterations in mentalizing processes have a high clinical relevance, since they occur before the onset of the disorder [17], persist over its course [15,18,19] and are linked to the patients’ outcome [20]. Potential dysfunctions within the neural network that underlies these social-cognitive alterations are, however, unknown. Therefore, we will leverage on advances in network neuroscience by using an analysis-scheme that determines each subject’s most likely network architecture while processing the task. Afterwards, the subject-wise networks will be condensed at the group-level to compare topological network alterations between patients with schizophrenia and healthy controls.

Previous neuroimaging studies revealed a set of core brain regions, which show enhanced functional activation during the inference of other people’s mental states. This suggests that the right temporo-parietal junction (rTPJ), the left TPJ (lTPJ), the medial prefrontal cortex (mPFC) and the precuneus/posterior cingulate cortex (PCC) are critical nodes of a network underlying mentalizing processes [21,22]. Previous research has frequently referred to the parallel activation of these brain regions as a network. However, the examination of the regions’ joint neural interactions during the mentalizing process is so far missing. Network approaches can thereby provide novel insight into the mutual influence of different brain regions within the network community. This can pave the way for a mechanistic understanding of mentalizing processes and potential alterations in psychiatric disorders, such as schizophrenia, to facilitate diagnostic procedures and treatment stratification.

Functionally, the regions within the mentalizing network have been found to be associated with social cognition and unconstrained thinking [23,24], leading to the assumption that the network might be relevant for a similar set of self – other computations [25,26]. This idea has been put forward by within-subject analysis, showing that the same regions are relevant for the processing of mental states and default mode cognition, which probably has been shaped by our species’ social nature [27]. Meta-analytical evidence further suggests that the midline-structures within the mentalizing network (dmPFC and PCC) are critical for social cognitive processes in general and self-other-related processing in particular. The bilateral TPJ on the other hand has been associated particularly with mentalizing processes and self-other distinction [28], which is why the fluent integration between these higher-order cognitions are likely fundamental for the emergence of complex social representations.

Clinical studies which have examined the neural basis of altered mentalizing processes in patients with schizophrenia found reduced activity in the core regions of the mentalizing network, such as the rTPJ [29,30,31], the lTPJ [30,31], the mPFC [30,31,32,33] and the precuneus/PCC [30, 34] during mentalizing tasks, indicating marked dysfunctions in regions relevant for mental states inferences. However, some studies found enhanced activation of these mentalizing brain regions in patients with positive symptoms, which could indicate a deficit in integrating self and other representations [32,34] or an ‘over-attribution’ of intentions [31] during mentalizing. A recent meta-analysis systemized these findings with converging evidence of functional activation alterations in two core regions of the mentalizing network, the TPJ and the mPFC, in patients with schizophrenia [35]. Although brain regions relevant for mentalizing processes have been consistently found to show altered neural responses in patients, the functional integration between them is, so far, less clear. This, however, is of high importance given that pathological states in the disorder can be understood as a dysfunctional integration of different cognitive processes [6,36], such as the attribution of an internal generated speech to an external source [7]. In this sense, it has been suggested that an online coding process, a dynamic information integration between distributed neural regions, by consistently adjusting edge strengths, i.e., their connectivity [37], might be altered in the disorder [6]. In this study, we empirically evaluate this hypothesis by examining, for the first time, specific network indices of the mentalizing network by means of graph theoretical methodologies [38,39]. In the terminology of graph theory, brain organization is considered as a network consisting of interconnections (edges) between spatially distributed neuronal populations (nodes). Accordingly, brain regions can be considered as hubs, which influence other brain regions within the network by their out-going connectivity (out-degree, the proportion of times a specific edge is present) or get influenced by other brain regions within the community (in-degree, the proportion of times a specific edge is present). Hubs are considered as important features of brain networks, given they distribute and integrate information within the community in a powerful way [40]. Their functional relevance increases with a higher amount of network connections. Hence, the elimination of highly connected brain regions can impact the global network function and might be associated with alterations in mental states. Therefore, we will describe potential integration deficits at a global (in-degree, out-degree) and a local level (edge-connectivity) to precisely capture the mentalizing network in patients with schizophrenia.

We used a network discovery approach [41] which allows to determine effective connectivity, i.e. the causal influence a brain region exerts over another within a functional network [42]. We examined this by means of an interactive mentalizing task, a social decision-making game, in which the participants need to infer the intentions of the playing partners to improve interpersonal decision-making [43]. The functional activity analysis was aligned with a meta-analytical derived functional mask of previous mentalizing studies [44] in order to define the core nodes of the network based on meta-analytical information. Then, we optimized each participant’s individual mentalizing network structure via dynamic causal modelling (DCM) [39,45,46] to identify the most likely functional mentalizing network architecture of each participant during the mentalizing task. Between-group comparisons focused on network indices such as the in-degree and out-degree of each hub and edge-connectivity between the regions (Fig. 1). Based on previous functional activation studies in patients with schizophrenia [31,32,34] and recent meta-analytical evidence [35], we expected to find integration deficits, particularly between the prefrontal cortex and temporo-parietal regions in patients during the mentalizing process.

**Fig. 1.**
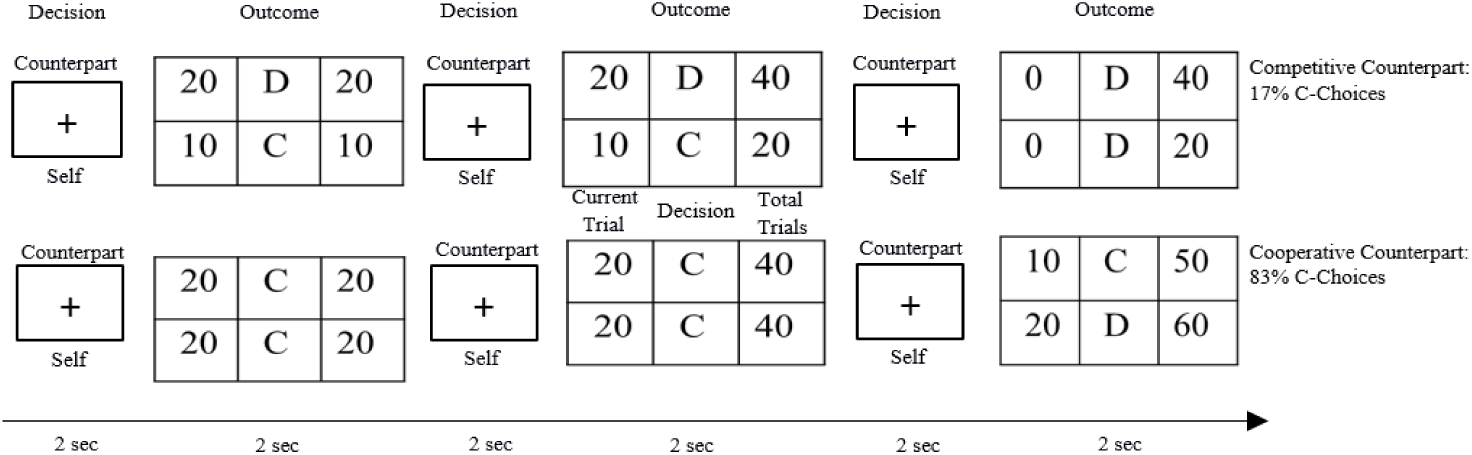
Examples of the interaction sequences with the intentionally different counterparts. Each interaction block contained six decision phases in which the participants had to decide to either defect (D) or cooperate (C) with the counterpart. Afterwards the partners’ payoffs were presented. The COMP and COOP playing partner decided in 5 out of 6 decisions per block with either the D-or C-choice.

## 2. Methods

### 2.1 Participants and Inclusion Criteria

A total of 31 (10 females) patients with a diagnosis of schizophrenia or schizoaffective disorder (ICD-10: F20 or F25) in the range between 18 and 53 years of age (M±STD=31.29±8.53) and 20 healthy controls (10 females) between 23 and 45 years of age (M±STD=29.55±6.17) were included in the analysis (for further sample characteristic see the results section). Both groups participated in a previous study, with a different research question [43]. A structured clinical interview for DSM-IV [47] was used to confirm the patients’ diagnosis and to exclude any psychiatric disorder in healthy controls. The severity of patients’ symptoms during the week before they participated in the study was assessed with the Scale for the Assessment of Positive (SAPS, [48]) and Negative Symptoms (SANS, [49]). Participants were only included by sufficient fMRI data quality (details in fMRI data preprocessing) and sufficient task engagement (not more than 25% misses of behavioral responses during the task). According to these criteria 9 patients (3 data-quality, 6 task-engagement) were excluded out of the original sample of 40 patients who participated in the study. All participants had normal or corrected-to-normal vision, reported no history of neurological disorders and were right-handed [50]. The authors assert that all procedures contributing to this work comply with the ethical standards of the relevant national and institutional committees on human experimentation and with the Helsinki Declaration. The study was approved by the local Ethics Committee at Philipps-University Marburg and all participants gave written informed consent prior to the experiment.

### 2.2 Experimental task

Before the experiment, participants were instructed that they will interact with three real playing partners in a social decision-making game in which the payoff depends on their own and the counterpart’s decisions (iterated prisoner’s dilemma game, for details: [43, 51, Fig.1]) this was practiced beforehand in a trainings session. In fact, the participants interacted with fictive playing partners who pursued a stable strategy either competitively (defecting in 83.3% of trials), cooperatively (cooperating in 83.3% of trials) or randomly (defecting/cooperating in 50% of trials respectively). The counterbalanced interaction blocks started with the presentation of a picture and a name of the current playing partner (3.5±1.25s), followed by a decision phase (2s) during which the participants had to decide to press the left (cooperate) or the right (defect) button, followed by the presentation (2s) of the decision and outcome for both players determined by the payoff matrix [43,51, Fig.1]. Participants interacted with each counterpart in alternate order in 7 interaction blocks, each containing 6 decisions. Because the individual outcome is determined by the participant’s and the counterpart’s decision (Fig.1) an interdependence between both interaction partners exists. Accordingly, the subjects can accumulate evidence about the playing partners’ intentions during the experiment and iteratively adapt their strategies accordingly. A condition in which the picture of the playing partner was replaced by a red cross and the hint that due to technical reasons no playing partner is connected, acted as control condition. During this condition, the participants had to alternately press the right and the left button with no outcomes being presented, i.e. no mentalizing processes were necessary. The control condition contained exactly the same sequences as the task conditions. The only difference between both conditions is that the outcome numbers are replaced by a hash in the control condition.

### 2.3 Behavioral Analysis

In order to examine differences in representing the different counterparts’ intentions during the task, we examined between-group differences in behavioral adaptation to the different counterparts (‘Dynamic-Tom’: task-dependent mentalizing index, which is the difference between defective decisions in the competitive vs. cooperative condition), according to our previous approach (see [43]). Additionally, we examined connectivity indices and mentalizing abilities with a task-independent measurement, the Reading the Mind in the Eyes Task (RMET), which we managed to collect for 33 participants.

### 2.4 fMRI data analysis

#### 2.4.1 fMRI data acquisition

All images were acquired using a Siemens 3-Tesla Trio, A Tim scanner with a 12-channel head matrix receive coil. Functional images were acquired using a T2* weighted single shot echo planar imaging (EPI) sequence (parallel imaging factor of 2 (GRAPPA), TE=30ms, TR=2000ms, flip angle 90°, slice thickness 3.6mm, matrix 64×64, in-plane resolution 3×3mm^2^, bandwidth 2232Hz/pixel, EPI factor of 64 and an echo spacing of 0.51ms). Data from 33 transversal slices oriented to the AC–PC line were gathered in descending order.

#### 2.4.2 fMRI data preprocessing

According to our previous approach [43] functional data preprocessing was performed using SPM8 (http://www.fil.ion.ucl.ac.uk/spm) implemented in MATLAB13a (MathWorks, MA). The first five volumes of each functional run were discarded from the analysis to account for T1 equilibration effects. Functional data were realigned and unwarped, corrected for slice timing, spatially normalized onto a common brain space (Montreal Neurological Institute, MNI) and spatially smoothed using a Gaussian filter with a 8mm full-width half maximum (FWHM). Prior to the analysis the fMRI-data was controlled for quality constraints. Participants were only included when less than 5% of the functional images were detected as outliers by an automated quality assurance protocol (Artifact Detection toolbox, https://www.nitrc.org/projects/artifact_detect/), based on massive head motion (> 2mm) or aberrant signal intensity (global-signal z-value exceeded a threshold of 9) criteria.

#### 2.4.3 fMRI data analysis

For each participant a general linear model (GLM) was calculated in which the decision and the outcome phase (4sec) were convolved with a hemodynamic response function (HRF) and implemented in the GLM with the presentation of the playing partners’ pictures and the 6 - realignment parameters (rigid body) as nuisance regressors. The high-pass filter was adapted to the experimental design and set to 284-sec cut-off period.

#### 2.4.4 Specification of dynamic causal models (DCM)

We used DCM12 as implemented in SPM12 (http://www.fil.ion.ucl.ac.uk/spm) to determine effective connectivity in the mentalizing network according to a network discovery approach [41]. In general, DCM allows to examine the causal influence one brain region exerts over another and how this coupling is influenced by changes in the experimental context [42]. We used the optimization procedure for post-hoc inferences [45,46] to discover each participant’s most likely network architecture, indicated by the greatest conditional probability in comparison to alternative networks [41]. Therefore, a full bilinear DCM was specified, subject-wise estimated and the most likely model selected by the Bayesian model evidence [45,52]. Individual network models (existence of connections between the regions) were used to set up the between-group comparisons, which focused on each node’s in-degree (the sum of present forward-connections in each subject of a group), out-degree (the sum of present backward-connections in each subject of a group) and edge-connectivity, i.e. the presence of an effective connection between the specific network nodes. Regions that were included in the dynamic causal modeling had to fulfill specific criteria in order to analyze commensurable functional areas [53]. Therefore, the region selection was guided by the activation peak of mentalizing brain regions found in the main effect (TASK>CONT) inclusively masked by a meta-analytically derived functional mask, covering brain regions relevant for mentalizing processes from 140 previous mentalizing studies (neurosynth; [44]). This strategy allowed us to limit our analysis to the mentalizing network found in previous studies of mentalizing processes. The analysis revealed activation in the core regions (the rTPJ, the lTPJ, the mPFC and the precuneus) of the mentalizing network across both groups (corrected for multiple comparisons, FWE, p<.05) which guided the extraction of each Volume-of-Interest’s (VOI) first eigenvariate of the regions (details in the results section). Each region’s first eigenvariate was extracted at a single subject level from all suprathreshold voxels (*p*=.05, uncorrected, main effect) of a 6mm sphere. The sphere was centered on the maximum of the local activation foci within 12mm search radius centered on the activation foci of the group analysis, adjusted for the effect of interest. Five patients and two controls showed no suprathreshold activation in one or more of the extracted mentalizing brain regions and where therefore excluded from the DCM analysis leading to a DCM sample of SZ: *n*=26 and HC: *n*=18. For the DCM analysis a novel first-level general linear model (GLM) was specified, which contained two regressors, one for the exogenous input (B-matrix), specified by the mentalizing conditions and the driving input (C-Matrix) specified by the entire task. Endogenous connections (A-matrix) were fully coupled between all four brain regions and the subject-wise connections (present or not) were the main objective of statistical analysis. Based on previous DCM-studies reporting group differences between patients with schizophrenia and healthy control subjects (N<20, e.g. 52, 54) we expected, that our fMRI sample size is sufficient to reveal the expected effects. Between-group analyses were conducted with Bonferroni-corrected (12 edge-connectivity, 8 in-/out-degree comparisons) chi-squared tests and Mann-Whitney U tests and considered as significant at α*<*.0025. Furthermore, we report results as trends when the analyses were significant at an uncorrected threshold of α*<*.05. Clinical characteristics were examined in patients with and without an absence of an edge-wise connection (please see supplementary material).

## 3. Results

### 3.1 Sociodemographic and Clinical Characteristics

No significant age differences existed between the groups (t(49)=.788, p=.434). Educational level (*χ*^2^=7.65, *p*=.054) and gender (*χ*^2^=2.57, *p*=.107) differed slightly between the groups, which is why both variables were added as covariates in the functional activation analysis. The patient group had a mean SAPS score of 12.35±10.57 (*M*±*STD*, range:0-36) and a SANS score of 10.35±11.74 (*M*±*STD*, range:0-39) leading to a global score of 22.70±19.44 (*M*±*STD*, range:0-73). The duration of illness was 9.35±8.40 years (*M*±*STD*, range:0–30 years). Twenty-four patients received stable doses of atypical antipsychotic medication (M*±ST*D=387.89±451.74mg/day Chlorpromazine equivalent [55], whereas seven patients did not receive any antipsychotic medication.

### 3.2 Differences in Adaptive Social Behavior during the Task

Healthy controls showed a higher adaptation to the intentionally different counterparts during the task, as significant between-group differences in the Dynamic ToM value suggest, t(49)=2.35, p=.023, HC: M±STD=5.5±6.48, SZ: M±STD=1.39±5.85.

### 3.3 Main effect TASK > CONT (Masked by the Mentalizing Network)

The main effect of mentalizing against the control condition (masked by the meta-analytically derived functional mentalizing mask) revealed enhanced activation in the core regions of the mentalizing network (Fig.1) such as the rTPJ (x= 60 y=-46 z=28, t=5.81, FWEp<.001, k=404), the lTPJ (x=-52 y=-50 z=36, t = 8.74, FWEp<.001, k=20), the precuneus (x=2 y=-64 z=46, t = 7.01, FWEp<.001, k=33) and the dmPFC (x=6 y=38 z=48, t=8.74, FWEp<.001, k=110) across groups.

### 3.4 Network Characteristics

Network analysis indicated that in patients with schizophrenia the dmPFC received overall lower input from other brain regions within the mentalizing network compared to healthy controls, U=98.0, p=.001 (Tab.1). Further edge-wise analysis indicates a significantly lower proportion of bilateral rTPJ to the dmPFC connections, χ2=10.47, p=.001 and of the lTPJ to the dmPFC, χ2=10.18, p=.001 in patients with schizophrenia compared to healthy controls (Tab.2, Fig. 2). Reduced overall out-going network connections of the dmPFC (U=132.0, p=.011), the lTPJ (U=119.0, p=.004) and the rTPJ (U=143.0, p=.023) in patients didn’t reach the Bonferroni corrected threshold (α<.0025). The same applies to the reduced lTPJ in-going connections from other brain regions within the network in patients with schizophrenia (U=152.5, p=.043) (Tab.2).

**Tab. 1.** In-degree and out-degree of each network hub

| Direction/Region | Schizophrenia<br>*M (STD)* | Controls<br>*M (STD)* | Mann-Whitney-U | *Z* | *p*-value |
| --- | --- | --- | --- | --- | --- |
| In_ITPJ | 0.5128 (0.329) | 0.7222 (0.366) | 152.5 | -2.026 | .043 |
| In_rTPJ | 0.6026 (0.377) | 0.7593 (0.358) | 173.5 | -1.536 | .125 |
| In_PREC | 0.5897 (0.380) | 0.7222 (0.357) | 185.0 | -1.235 | .217 |
| In_mPFC | 0.3462 (0.358) | 0.7593 (0.339) | 98.0 | -3.368 | .001* |
| Out_ITPJ | 0.5000 (0.316) | 0.7778 (0.228) | 119.0 | -2.884 | .004 |
| Out_rTPJ | 0.4872 (0.301) | 0.7037 (0.300) | 143.0 | -2.278 | .023 |
| Out_PREC | 0.5641 (0.336) | 0.7222 (0.307) | 171.0 | -1.576 | .115 |
| Out_mPFC | 0.5000 (0.341) | 0.7593 (0.375) | 132.0 | -2.538 | .011 |

In = Number of effective connections a network node receives from other brain regions. Out = Number of effective connections a network node integrates into other brain regions. \* Significant Bonferroni corrected between-group differences are indicated.

**Tab. 2.** Edge-wise effective connectivity between the brain regions and group differences

| From region → region | Proportion Present<br>Schizophrenia | Proportion Present<br>Controls | $\chi^2$ | *p*-value |
| --- | --- | --- | --- | --- |
| ITPJ → mPFC | 0.3462 | 0.8333 | 10.182 | .001* |
| ITPJ → rTPJ | 0.5769 | 0.7778 | 1.91 | .167 |
| ITPJ → PREC | 0.5769 | 0.7222 | .970 | .325 |
| rTPJ → mPFC | 0.2308 | 0.7222 | 10.471 | .001* |
| rTPJ → ITPJ | 0.4231 | 0.7222 | 0.970 | .325 |
| rTPJ → PREC | 0.6538 | 0.6667 | .008 | .930 |
| PREC → mPFC | 0.4615 | 0.7222 | 2.946 | .086 |
| PREC → rTPJ | 0.7308 | 0.7778 | .125 | .723 |
| PREC → ITPJ | 0.5000 | 0.6667 | 1.204 | .272 |
| mPFC → rTPJ | 0.5000 | 0.7222 | 2.713 | .140 |
| mPFC → ITPJ | 0.4615 | 0.7778 | 4.40 | .036 |
| mPFC → PREC | 0.5385 | 0.7778 | 2.632 | .105 |

Effective endogenous connectivity between two brain regions (edge-connectivity) and their direction (indicated by the arrow) between the groups. \* Significant Bonferroni corrected between-group differences are indicated.

**Fig. 2.**
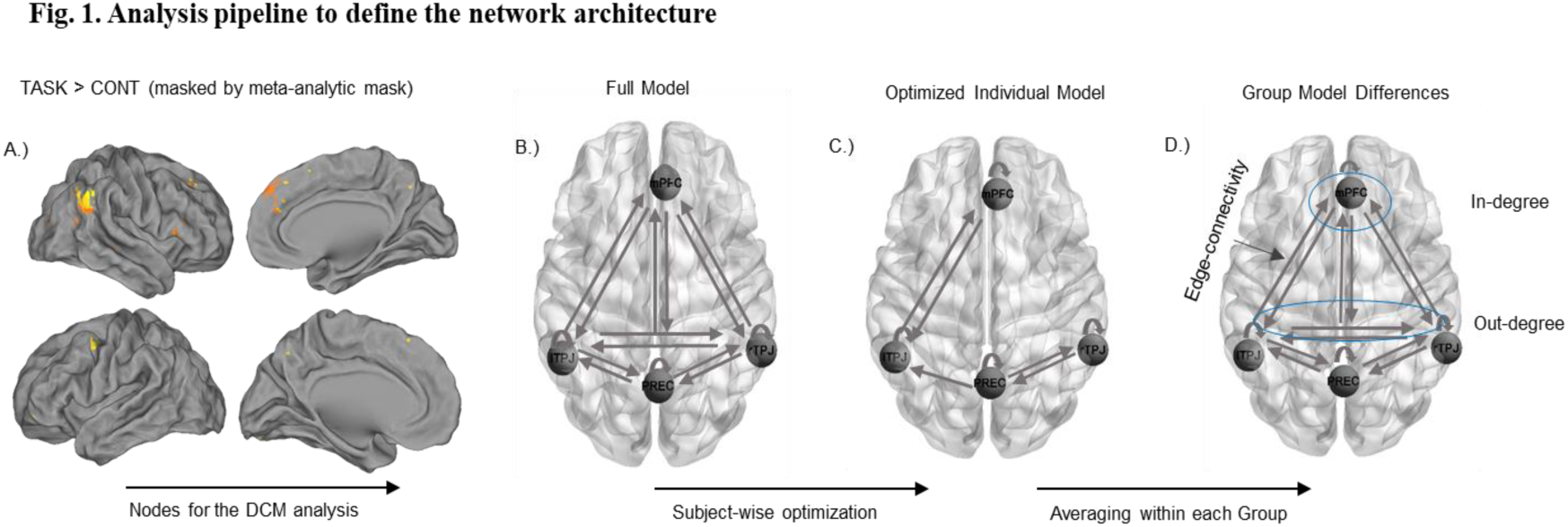
A) The main effect of mentalizing conditions was masked by a meta-analytic mask (140 mentalizing studies) to align the current study with evidence from further mentalizing studies and guided the selection of the nodes for the Dynamic Causal Model (DCM) B) Example of a fully connected network including the precuneus (PREC), the left and right temporo-parietal junction (lTPJ,rTPJ) and the dorsal medial prefrontal gyrus (mPFC) was optimized for each subject C) Example of an optimized model, which indicates the most likely network structure D) Edge-connectivity, in and out-degree per network node were calculated for each group and compared.

**Fig. 3.**
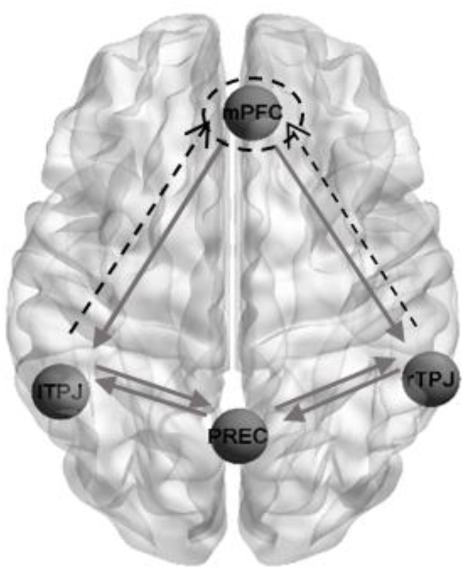
Differences in the network architecture between the groups, point on significantly reduced effective connections from the right and left temporo-parietal junction (rTPJ,lTPJ) to the dorsal medial prefrontal cortex (dmPFC, dashed lines) in patients with schizophrenia compared to healthy controls. Furthermore, in patients with schizophrenia the dmPFC received less frequently input from other brain regions in the mentalizing network (dashed circle).

### 3.5 Network Characteristics and Task Dependent / Independent ToM-Indices

Additionally, we examined potential associations between the Reading the Mind in the Eyes Task (RMET), as a task-independent index for mentalizing abilities and network characteristics. Notably, no significant between-group differences were observed for test scores in RMET (t(31)=1.69, p=.101; HC: M=26.56±3.56, Range: 33-21; SZ M=24.53±3.37, Range: 30-19). However, exploratory analyses revealed that the network characteristic is associated with task-independent and task-dependent indices of mentalizing, as suggested by significant higher RMET values in patients with vs. without an edge-connection in every network node (lTPJ-rTPJ (t(31)=3.04, p=.005), MPFC-lTPJ (t(31)=2.42, p=.022), rTPJ-PREC (t(31)=2.44, p=.021)). Given that these analyses were not corrected for multiple comparisons, they should be considered as exploratory. Along this line, the subjects’ understanding of the counterpart’s intentions (Dymanic ToM) during the task correlated positively with the entire network structure (richness of edge-connections within the network; r=.257, p=.046, one-tailed). Hence, internal (task-dependent) and external (task-independent) associations indicate an association between the ToM network structure and mentalizing processes. Given that the network characteristic and ToM-indices analysis are not corrected for multiple comparisons, they should be considered as exploratory.

### 3.6 Network Alterations and Medication

In order to exclude that network alterations are influenced by medication, we examined a potential relation between-group network differences and the antipsychotic dose in patients (chlorpromazine equivalent). These analyses showed no significant associations between medication and the in-degree of the dmPFC in patients (*r*=-.068, *p*=.74), nor a higher dose in patients with vs without edge-wise connectivity between the rTPJ - dmPFC (*t*(24)=.66, *p*=.51) and the lTPJ - dmPFC (*t*(24)=-.28, *p*=.78). Additional control analysis can be found in the supplementary material.

## 4. Discussion

In this study we characterized the effective connectivity between brain regions forming the mentalizing network in patients with schizophrenia for the first time. The DCM network discovery approach indicates a sparser network architecture in patients and highlights the dmPFC as a disconnected network hub, since the region’s activation changes are less influenced by neural activity changes of the right and the left TPJ. These findings provide first evidence that alterations within the core mentalizing network arise due to disconnections between at least three core regions of mentalizing processes in patients with schizophrenia and pin-point dysfunctions particularly of the forward TPJ connectivity.

### 4.1 A network perspective can facilitate the understanding of alterations in patients’ mentalizing processes

The examination of functional networks can facilitate the understanding of phenotypical alterations in mental disorders [1,2]. Similarly, alterations in brain networks have been found in depression [52] and autism [56]. Schizophrenia has been associated with disturbances in the interaction of distant brain regions, indicating that neural ‘disconnections’ contribute to the disorder’s pathophysiology [3,4,6] underlined by findings of disruptions in functional networks in childhood-onset schizophrenia [57]. One characteristic epiphenomenon of schizophrenia is impaired social functioning [58], which is linked to impairments in mentalizing processes [15] and might be affected by disrupted information processing between distant brain regions [59]. By investigating effective connectivity during a social decision-making game, the current DCM analysis [41] allowed us to pin-point how brain regions within the mentalizing network influence each other and whether their forward or backward effective connectivity is altered under restricted task conditions.

Our findings reveal that in patients with schizophrenia the dmPFC arises as a disconnected network hub, due to a dysfunctional effective forward connectivity from the bilateral TPJ, indicating an impaired information integration within the mentalizing network. Further analyses suggest that the integration between the regions within the network is associated with enhanced general mentalizing abilities, indicated by significant associations with the Reading the Mind in the Eyes task, a task-independent index for mentalizing abilities. On a system level our findings indicate that the connection intactness within the entire network is linked with an enhanced ability to generate representations of different interaction partners during the task, pointing to a link between the plethora of interconnections within the mentalizing network and beneficial social behavior during the task.

The question arises how these network dysfunctions translate into alterations in the mentalizing process in schizophrenia. Previous research suggests that the brain regions in the mentalizing network are implicated in mnemonic, prospective and default mode processes, indicating a common core function, such as the projection into another constructed perspective beyond the current environment [60]. Along this line, it is assumed that some forms of mentalizing processes draw on the own mental experience to model other people’s mental states [60,61], probably by guiding mentalizing processes through self-knowledge [62], which has been associated with an intact dmPFC function [60,62,63]. Accordingly, it has been found that the processing of introspective information and inference processes of other people’s mental states converge in the dmPFC [64,65], which might be related to the region’s role in social interactions [65]. Furthermore, it has been suggested that the dmPFC is implicated in the generation of higher-order, particularly sensory-independent social cognitive processes, which is reflected in the region’s interconnection with associative and integrative brain regions, such as the TPJ [67]. Hence, the current findings of reduced neural integration between the TPJ and the dmPFC in patients might indicate a diminished incorporation of other peoples’ mental states in higher-order, stimulus-independent processing modes.

Previous findings suggest that in patients with schizophrenia impairments in generating mental representations of others’ mental states are associated with dysfunctions in the functional connectivity of the rTPJ to brain regions in the temporal lobe [43]. This points to alterations in the liaison between mentalizing and memory processes [51] which is in line with behavioral findings [68]. Notably, these previous results might be associated with the present findings of disconnections in patients’ mentalizing network. During the experiment, the subjects can accumulate evidence about the playing partners’ intentions and iteratively adapt their strategies, leading to a continuous updating of their mental representations, particularly in the TPJ [51]. Previously it has been shown that impaired TPJ updating results from a dysfunctional integration of memory information from previous experiences in the hippocampus and other temporal lobe regions [43]. This might be associated with the TPJ’s impaired effective forward connectivity and could lead to an incomplete integration of mental models of others’ intentions into dmPFC processes in patients with schizophrenia. Since the TPJ [69] and the dmPFC [67] are neuroanatomically connected with brain regions relevant for memory processes, a consequent analysis how this cognitive mechanism relies on episodic memory information requires further examination. So far, there is evidence from neural [43] and behavioral studies [68] that episodic memory impairments account for deficits in mental state inferences in patients with schizophrenia. Furthermore, our findings of a disconnected frontal hub in the mentalizing network is in line with previous findings, which indicate that the frontal cortex receives less information from distant brain regions in patients with schizophrenia [13, 70] and begs the question whether these findings extend to alterations in further cognitive and symptomatic domains.

### 4.2 Dysconnectivity between the lTPJ and the dmPFC has been associated with psychosis risk

Remarkably, a genome-wide association study has previously found that healthy individuals with a high risk for psychosis show reduced connectivity between the dmPFC and the lTPJ during a mentalizing task [71]. Taking our and previous findings together, the dysfunctional connectivity between the TPJ and the dmPFC could be a valuable biomarker that might be present before the onset of the disorder and can also be found in patients with a diagnosis. Importantly, the lTPJ has been found to be associated with hallucinatory experiences. Accordingly, when direct electrical stimulation is applied to the region the emergence of a visual hallucination has been reported in a non-psychiatric person [72]. Vice versa, a reduction of auditory hallucinations has been reported following a cathodic tDCS-application in patients with schizophrenia [73], indicating that the TPJ might be a critical target region in the pathophysiology of schizophrenia. Evidence from tDCS-studies suggest that the rTPJ is associated with an abstract process of taking a perspective to view another person’s mind [74,75,76], and that the dmPFC facilitates the integration of these processes into a higher order representation of a social constellation [65,74]. Future brain-stimulation studies have therefore to show whether the TPJ modulation can facilitate the dmPFC’s integration and thereby improve mentalizing abilities in patients with schizophrenia.

### 4.3 Limitations and future perspectives

The findings should be interpreted in the light of some limitations. First, DCM is a hypothesis driven analysis strategy in which the selection of brain regions is conducted a-priori. Therefore, it should be noted that alterations in the complex cognitions which occur during social interactions, such as mentalizing, might be further facilitated by secondary cognitive processes (e.g. memory, attention or self-processing), or might be considered their complex conglomerate. Accordingly, recent research suggests that mentalizing can be considered a multidimensional construct, subsuming and integrating different sets of cognitive mechanisms [22,77,78], which might be particularly relevant for patients with schizophrenia [43]. Along this line, it should be considered that in total seven participants failed to engage specific nodes within the specified network, which is a common issue in DCM studies [79]. Although several reasons can account for an individual’s missing activation of a specific brain region [80], it might be that the excluded participants used an alternative strategy to perform the task and thus recruit different brain regions. Future studies should therefore investigate whether the joint interaction of other brain regions or networks with the mentalizing network can compensate or potentially account for the topological alterations we have found in this study. Another emerging question is how our findings impact real-life interactions in healthy participants and patients with schizophrenia. Therefore, future studies might assess how the amount and the quality of an individual’s daily social life shapes the neurobiology of the mentalizing network. Hence, it could be clinically relevant to examine whether the outcome of psychological (group) interventions, which can be considered as structured and goal-oriented social interactions, are modulated by the intactness of the neural substrates underlying social interactions or whether therapeutic processes can lead to a reorganization of previously altered network structures.

However, by interpreting the findings it should be considered that, in the current task, mainly the cognitive aspects (as opposed to affective components) of the mentalizing process are assessed. Hence, future studies might examine whether alterations in the dmPFC-integration extend to alterations in affective mentalizing processes in patients with schizophrenia. A further limitation of our study is the medication intake in patients. Although, we statistically could not detect a significant association between alterations in the mentalizing network and medication intake, it should be considered that the healthy and patients’ group did differ in this relevant variable. Accordingly, it should be considered that the duration of illness in our sample was quite variable. Further large-scale studies might therefore examine in specific sub-samples whether the duration of illness influences the mentalizing network architecture in patients with schizophrenia. These aspects should be kept in mind during the interpretation of the results. The aim of this study, however, is to examine alterations in the architecture of the core network for mentalizing processes in patients, which might be an initial starting point for therapeutic approaches.

## 5. Conclusion

The characterization of the mentalizing network in patients with schizophrenia revealed a sparser structure in patients, particularly due to a disconnected dmPFC, which is less driven by bilateral TPJ computations. Our findings highlight the relevance of the connectivity profile of the TPJ for social cognitive processes in schizophrenia, rendering it to a potential target region for neurostimulation.

## Acknowledgments

This work was supported by the Core Facility Brain Imaging, Faculty of Medicine, University of Marburg, Else Kröner-Fresenius-Stiftung (grant number: 2014_A136) and Deutsche Forschungsgemeinschaft (grant numbers: STR-1146/4-1, STR-1146/8-1, STR-1146/9-1, to BS)

